# Salient auditory stimuli evoke spatially segregated phasic and sustained neural responses in the human brain

**DOI:** 10.64898/2025.12.18.695315

**Authors:** Siddhartha Joshi, Mensure Polat, Daniel C. Chai, Sofia Pantis, Ritika Garg, Vivek P. Buch, Ashwin G. Ramayya

## Abstract

Salient sensory stimuli are known to evoke neural activations across distributed brain regions. However, the temporal dynamics of these responses over sub-second timescales remain poorly understood, in part due to limitations in the temporal resolution of non-invasive neuroimaging methods. We examined the spatiotemporal dynamics of neural activations evoked by salient sensory stimuli (rare sounds) using 1,194 widely distributed intracranial electrodes in 5 neurosurgical patients. Salient stimuli preferentially activated 263 of 1,194 electrodes (22%), with responses segregating into two largely distinct spatiotemporal patterns: (1) phasic activation in sensorimotor regions, and (2) sustained activation within the salience network. Cross-correlation analysis revealed that phasic sensorimotor activation preceded sustained salience network activation on a trial-by-trial basis. These findings support an updated view of salience processing in the human brain, revealing that salient stimuli evoke two sequential stages of neural activation—phasic sensorimotor responses followed by sustained salience network activity—rather than simultaneous widespread activation.

## Introduction

Salient sensory stimuli are those that easily stand out in the environment. For example, a sudden change in our sensory (visual or auditory) surroundings while driving re-orients our attention towards specific vehicular or pedestrian traffic, optimizes brain processes that detect and process danger, and generates a response towards safety. How are such salient sensory stimuli encoded in the human brain? Experimental studies have shown that the detection and encoding of salient information are key steps in neural processes that mediate adaptive behaviors key to survival, including attention and learning (*Watkins et al., 2007; Kerzel D and Schönhammer J., 2013; Wang et al., 2014; Keil et al., 2016; Liu et al., 2022; Novembre et al., 2024*). Theoretical explorations have highlighted the importance of salience detection and encoding for tracking statistical irregularities and generating prediction error signals, both of which are central components of widely used models of attention and learning (*Yu and Dayan, 2005; Cohen and Aston-Jones, 2005; Wilson et al., 2010; Nassar et al, 2010*).

Human functional magnetic resonance imaging (fMRI) experiments have shown that salient sensory stimuli, such as rare “oddball” sounds or visual targets, evoke neural activation in auditory or visual cortices (depending on the stimulus modality), and in regions such as dorsal anterior cingulate cortex and the anterior insula that are major hubs of the “salience network” (*Linden et al., 1999; Stevens et al., 2000*; Greicius et al., 2003; *Liebenthal et al., 2003; Seeley et al., 2007; Critchley et al., 2011; Beissner et al., 2013; Murphy et al, 2014; Seely, 2019*). Oddball stimuli reliably evoke event-related potentials, recorded using scalp electroencephalography (EEG) in human subjects, and the timing and size of these evoked potentials is correlated with oddball detection performance (*Hillyard et al., 1971; Parasuraman and Beatty, 1980; Murphy et al., 2011*). Interestingly, evoked potentials associated with surprising or cognitive aspects of salience tasks show distinct temporal dynamics, but the neural circuits in the human brain that give rise to these externally measured signals have remained largely unknown (*Näätänen et al, 1978; Javitt et al., 1994; Murphy et al, 2011; Camalier et al, 2019; Obara et al., 2023*).

Despite the rich spatial information from fMRI studies, their limited temporal resolution means that sub-second neural dynamics are poorly understood. In contrast, scalp electroencephalography (EEG) recordings have high temporal resolution but limited spatial resolution (*Brázdil et al., 2005; He et al., 2023*). Although a few human studies have explored salience processing using high spatiotemporal resolution intracranial EEG (iEEG) recordings, they are limited to selected regions of interest or analyses of low frequency components of the iEEG signal, which are difficult to interpret in terms of local neural spiking activity (*Paller et al, 1992; Zaehle et al, 2013; Citherlet et al, 2019, 2020; Ferraro et al., 2020; Mocchi et al, 2024*). Therefore, we have a poor understanding of the spatiotemporal dynamics of human brain neural activations evoked by salient sensory stimuli. This knowledge gap hinders validation of theoretical models which explain how the human brain encodes and transforms salient sensory signals to guide adaptive behavior.

Animal studies have provided a glimpse into the rich spatiotemporal dynamics of neural activation patterns in response to salient stimuli. For example, single unit studies in animal models have shown that the initial neural activation in response to salient sensory events is typically phasic (brief or transient) in a limited set of brain areas, including early sensory regions like inferior colliculus and primary auditory and visual cortices, and also in brainstem nuclei that are known to regulate sympathetic arousal (*Rajkowski et al., 1994; Swick et al., 1994; Fishman and Steinschneider, 2012; Joshi et al, 2016; Camalier et al, 2019; Gong et al., 2024*). Following this initial response, later activation is found in higher-order, associative regions including parietal and prefrontal cortices and the amygdala (*Ulanovsky et al., 2003; Chen et al., 2015; Takaura and Fujii, 2016; Camalier et al., 2019; Gong et al., 2024*). But whether these distinct neural dynamics in specific regions and networks are also found in the human brain remains largely unknown.

Motivated by these prior studies, we hypothesized that neural activations evoked by salient sensory stimuli unfold across multiple stages of spatiotemporal processing, rather than a simultaneous widespread activation. We tested this hypothesis by using intracranial electrophysiology (iEEG) in patients undergoing monitoring for medically refractory epilepsy as they performed a standard auditory oddball paradigm. We analyzed high frequency activity as a surrogate for neuronal spiking activity and performed unsupervised clustering to segregate electrodes based on their temporal dynamics. We found that salient stimuli evoke two sequential stages of neural activation—phasic sensorimotor responses followed by sustained salience network activity—rather than simultaneous widespread activation.

## Methods

### Participants

In this study, we recruited 5 patients with medically refractory epilepsy who underwent surgical implantation of intracranial electrodes for seizure localization. Patients volunteered to perform cognitive testing as part of our research study while they were admitted to the hospital and provided informed consent. These patients will hereby be referred to as “participants”. Our study was approved by the Stanford University Institutional Review Board. Clinical circumstances alone determined the number and placement of electrodes implanted. All ethical regulations relevant to human research participants were followed.

### Auditory oddball task

We used a standard auditory oddball task with contingency reversal (Figure 1A). The task was created and deployed using Python and PsychoPy running on a Dell laptop. In order to synchronize intracranial neural recordings with task events, timing signals were sent via a Labjack T4 device to the clinical EEG system. Each session started with verbal instructions given to the participant. We asked participants to maintain their gaze on a fixation cross displayed at the center of a laptop screen, keep count of the number of oddball stimuli, and report this number at the end of each block of trials. We consider the oddball tone to be salient both because of its statistical rarity and because subjects were tasked with counting only these tones, i.e. the tone was perceptually salient and also relevant for the task. These instructions were also displayed as text on the laptop screen prior to the appearance of the fixation cross. In each session, we ran two blocks of 100 trials each, switching the oddball tone in the second block. This switch was used to ensure that enhanced responses to the oddball tone were driven by its statistical rarity rather than by neural selectivity for a particular tone frequency. The stimulus timing and intertrial interval were both randomized to diminish stimulus timing predictability with interstimulus times varying between 2.5 and 4 sec.

**Figure 1.**
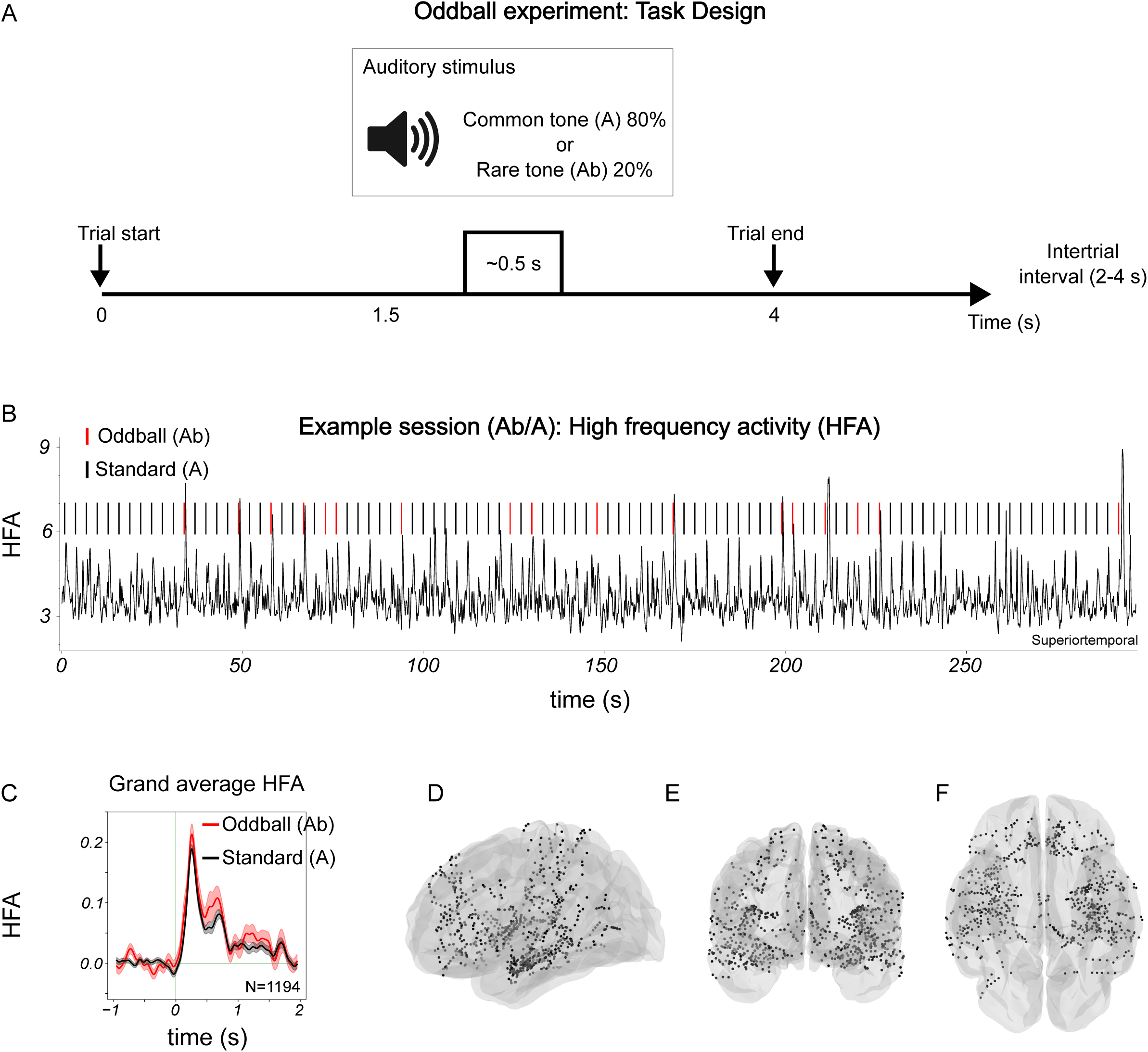
Task, examples and grand average responses to stimuli. (A) Oddball task design. (B) Example HFA responses during one oddball experiment session. (C) Grand average responses from all recording sites (N=1194). (D-F) Locations of all recording sites.

### Intracranial neural recordings

Intracranial EEG was recorded via depth electrodes (Ad-Tech or PMT Corporation) using the Nihon Kohden recording system with signals sampled at 1kHz. Signals from contacts on a single electrode were converted to a bipolar montage by taking the difference of signals on immediately adjacent contacts. The resulting bipolar signals were treated as new virtual electrodes (“electrode” hereafter) with location at the midpoint of each pair. We excluded electrodes with prominent line noise (60Hz), defined as greater spectral power in the 58-62Hz range than in the 18-22 Hz range, or electrodes that were disconnected (signal standard deviation=0).

### Anatomical localization of electrodes

Intracranial electrodes were identified manually on each post-operative CT scan and contact coordinates recorded based on the center of density of the radiodense contacts on thresholded images. To obtain contact locations in each patient’s native anatomic space as well as a common reference space (MNI coordinates), we registered the post-operative CT to the pre-operative MRI, and the MRI to the Montreal Neurological Institute (MNI) average brain. We assigned each electrode to various canonical intrinsic brain networks (“7 network model”) using a volumetric atlas in MNI coordinates (Yeo et al., 2011). We refer to the “ventral attention” network as the “salience” network, but otherwise use terminology as reported in the original study.

### Extracting high-frequency activity (HFA)

We extracted three seconds of iEEG data around each auditory stimulus presentation, one second preceding the stimulus and two seconds following it. This 3 second window defined one trial. We first used Welch’s method to estimate power spectral density between 0 and 500 Hz. We identified and rejected bad and disconnected electrodes as defined above (“<u>Intracranial neural recordings</u>”). We then extracted high-frequency spectral power for each trial using the continuous wavelet transform (PyWavelets: Lee et al., 2019). We used a Morlet wavelet with bandwidth parameter 1.5 and center frequency 1.0 to optimize the tradeoff between time and frequency resolution, and logarithmically spaced frequencies between 70 and 200 Hz. We then squared the wavelet convolution, log transformed the resulting power values, smoothed with a Gaussian filter (σ=30 ms) and then standardized each trial time-series by z-scoring using the mean and standard deviation from all trials in the block followed by subtracting the average activity just preceding each tone (baseline subtraction). Hereafter, we refer to this log-transformed, z-scored, baseline-corrected HFA as “HFA”.

### Analyzing responses to auditory stimuli

After examining individual trial HFA and per-electrode, trial averaged values, we defined a response window between 0.25 and 0.75 sec following each tone. For each trial, the HFA within this window was averaged to obtain the response to the tone. For each electrode, we used a non-parametric test to determine whether the oddball stimulus reliably evoked a larger response than the standard stimulus (p<0.05, Mann-Whitney U-test for *H_0_*: response to oddball tone > response to standard tone). These electrodes were defined as “responsive”. If the oddball tone evoked a response that was smaller than the standard tone response, we defined these as “suppressed” responses. The remaining electrodes were termed as “unresponsive”.

### Hierarchical clustering of neural populations based on response profiles

We used data-driven unsupervised clustering (agglomerative clustering, Scikit-learn) to group responsive electrodes based on activation patterns. Oddball tones typically evoked larger responses than standard tones. Therefore, we transformed the HFA to a uniform scale such that the range of values ranged from 0 – 1 prior to clustering since we wanted the clustering to identify only temporal response patterns without being biased by differences in response amplitudes. Once clusters were identified using the normalized HFA, per-cluster, tone-aligned responses were plotted using the regular HFA (as defined above).

### Cross-correlation analysis

We performed a cross-correlation analysis to assess the temporal relation between phasic and sustained clusters. For each trial, we averaged responses from all electrodes in each cluster to obtain a per-trial cluster-averaged response. Thus, for each participant, for each oddball tone, there was a single response for the phasic cluster and a single response for the sustained cluster. We also performed this computation for each standard tone. We then calculated the trial cross-correlation between phasic and sustained clusters separately for the oddball and standard tones. We estimated noise correlations using time windows that excluded tone-evoked responses. For each trial, we subtracted the noise correlation from each trial’s signal correlation at each lag (“corrected” signal correlation). We then averaged the corrected signal correlation across all oddball trials and standard tone trials separately. 95% confidence intervals were computed from the aggregate distributions of noise-subtracted time series across each pair of electrodes.

### Code

We wrote custom code scripts in Python for all data analysis. We used open-source scientific libraries and repositories including NumPy, SciPy, Scikit-learn, Pandas, MNE, and Nilearn. Experiment code was written in Python and PsychoPy. All code and data for this report are available at: https://github.com/RamayyaLab.

## Results

### Responsive electrodes

We found selective neural activations following oddball sounds in 22% of all electrodes (263 of 1,194 bipolar pairs across 5 participants). We measured neural activity using high-frequency activity (HFA, 70–200 Hz power), which provides a reliable surrogate of local (within ~3 mm) neural population spiking activity (Manning et al., 2009; Ray and Maunsell JH, 2011; Dubey and Ray, 2019; Figure 1A, D-F). Of these, 263 (22% of all electrodes) had reliably larger responses to the oddball than to the standard tone (“responsive” electrodes; example in Figure 1B; group average in Figure 2A; population data in Supplementary Figure 2A,B) while 848 (71% of all electrodes) showed no reliable effect of the oddball tone (“unresponsive electrodes”; Figure 2B). Neural activity at 83 electrodes (~7% of all electrodes) was suppressed below baseline by the oddball tone (Supplementary Figure 2D,E). These suppressive responses will be considered elsewhere. We found no electrodes that showed a reliably larger response for the standard relative to the oddball tone (Supplementary Figure 2G,H). Responsive electrodes were distributed among the sensorimotor, salience (ventral attention) and frontoparietal networks while unresponsive electrodes were in the default mode, limbic, and dorsal attention networks (Figure 2C,D,E,F).

**Figure 2.**
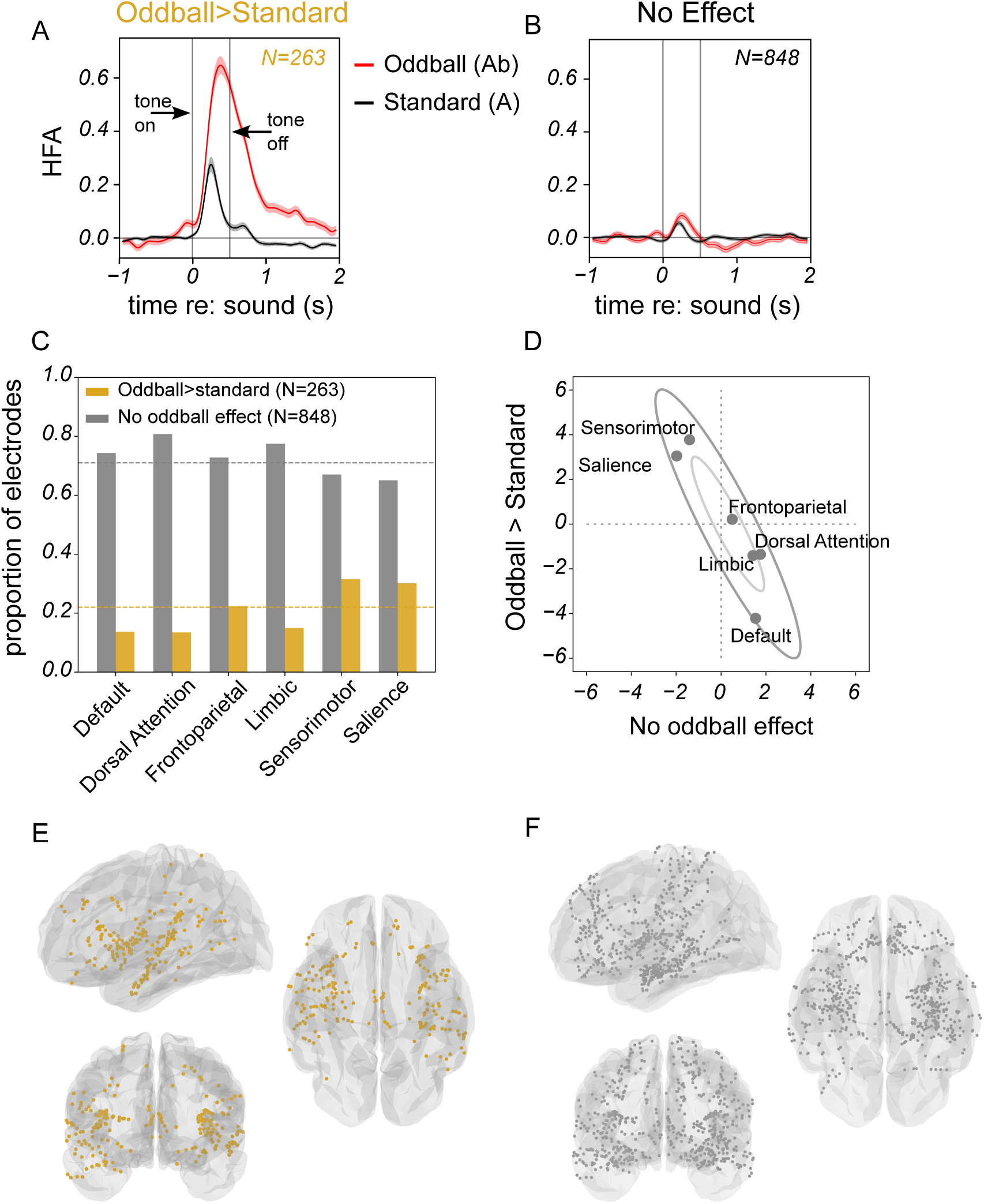
Responsive and unresponsive groups. (A) High frequency activity (HFA; mean +- sem) aligned to sound for oddball and standard stimuli, averaged over locations that showed reliable responses. (B) Same as (A) for non-responsive locations. Vertical gray lines indicate time during which tone was played. (C) Distribution of electrodes among standard brain networks. (D) Relative frequency of electrodes with phasic (ordinate) or sustained (abscissa) responses to the oddball stimulus (z-scores). Inner and outer ellipses are 1α and 2α confidence intervals derived from the joint distribution, respectively. (E,F) Anatomical locations for these groups of electrodes.

### Distinct temporal profiles of responsive electrodes revealed by clustering

We found that responsive electrodes showed two dominant temporal dynamics: phasic and sustained (Figure 2A). We used a hierarchical clustering method (Scikit-learn, AgglomerativeClustering) to segregate electrodes based on their temporal dynamics. We selected the clustering level that identified clusters that generalized across all participants, such that each participant had at least one electrode in each cluster (Ramayya et al., 2025). This approach revealed two clusters with distinct temporal profiles. The first cluster showed responses that peaked early (median [interquartile range]=0.27 [0.32 0.36] seconds; Figure 3A; Supplementary Figure 3A) and returned to baseline within 1 second following tone onset. Because of the relatively brief nature of these responses, we termed this cluster as “phasic”. The second cluster had relatively later peaks compared with the phasic cluster (0.48 [0.38, 0.59] seconds, Figure 3B; Supplementary Figure 3A) and responses that remained elevated for >1 second after tone onset. We termed this cluster as “sustained”.

**Figure 3.**
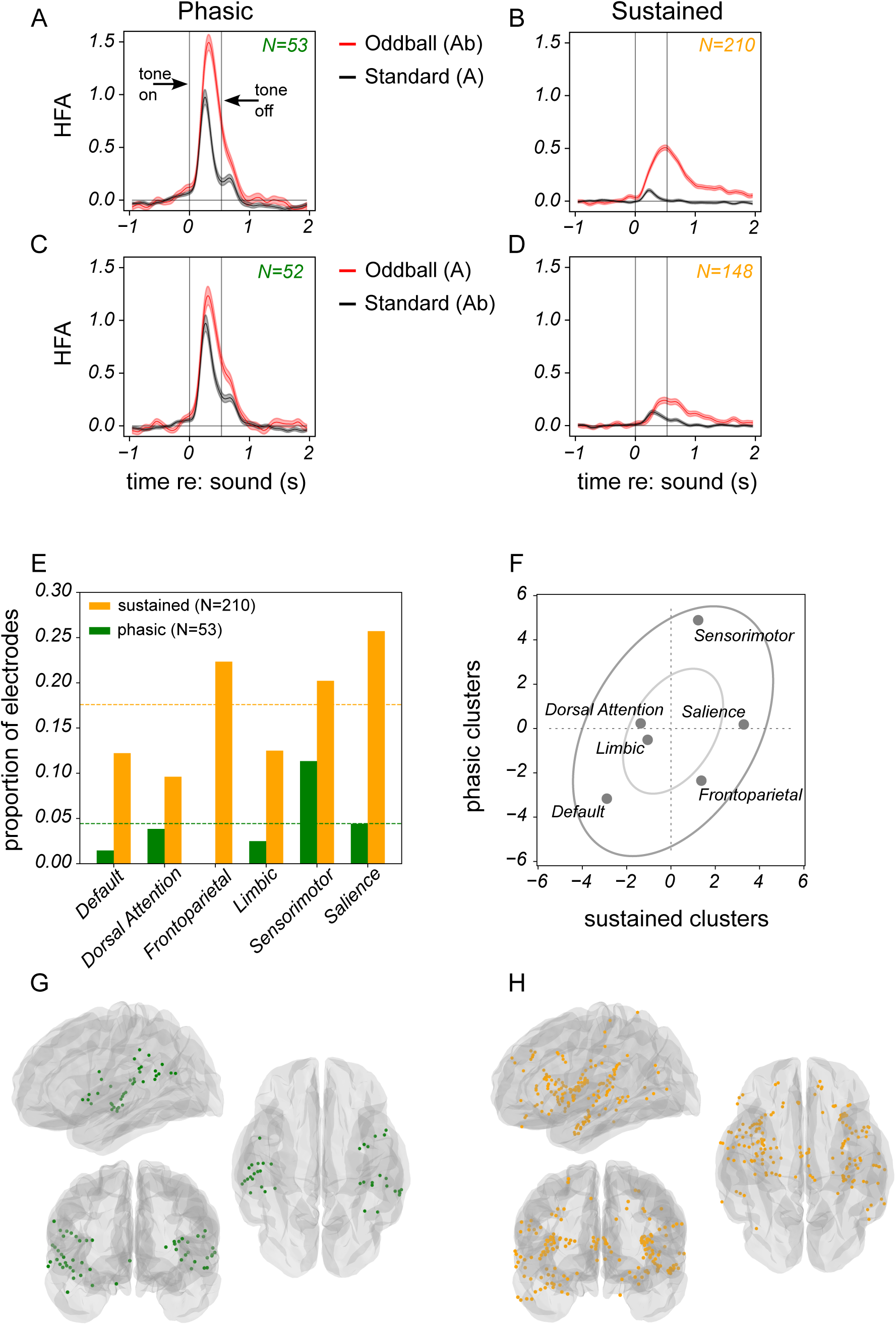
Distinct temporal profiles of responses to oddball stimuli are anatmomically segregated and are preserved with contingency reversal. (A) High frequency activity (HFA; mean +/- sem) aligned to sound for oddball and standard stimuli, for the phasic cluster. (B) Same as (A) for sustained cluster. Vertical gray lines indicate time during which tone was played. (E) Distribution of electrodes among standard brain networks. Horizontal, dahed lines indicate the proportion expected if the responses were distributed uniformly. (F) Relative frequency of electrodes with phasic (ordinate) or sustained (abscissa) responses to the oddball stimulus (z-scores). Inner and outer ellipses are 1α and 2α confidence intervals derived from the joint distribution, respectively. (G,H) Anatomical locations for these groups of electrodes.

### Anatomical locations of phasic and sustained clusters

Phasic and sustained responses were found brainwide but were distributed highly nonuniformly across the standard brain networks (Figure 3E; for both phasic and sustained: p < 0.005 (dof=0), chi^2^ statistic (phasic)=42.59, chi^2^ statistic (sustained)=23.15, chi-square test for *H_0_*: numbers of observed phasic and sustained sites are obtained by independent sampling from a uniform distribution of these responses across the brain). The two response distributions were also reliably different from each other (Figure 3E; p < 0.005, chi^2^ statistic=0.42, dof=5; chi-square contingency test for *H_0_*: pattern of distribution of phasic and sustained responses is similar across the standard brain networks). Phasic responses were dominated by sites identified as part of the sensorimotor network (Figure 3E,F,G), with ~74% (39/53) located in the superior temporal region, likely including auditory cortex (Yeo et al., 2011; Uddin et al., 2019). Sustained responses were found mainly at sites in the salience (or ventral attention; 64/210 or 30%) network, with smaller proportions in the frontoparietal (23/210 or 11%) and sensorimotor networks (57/210 or 27%; Figure 3E,F,H).

### Response profiles preserved after tone reversal

The two clusters preserved their response profiles following contingency reversal, i.e. after the tones associated with the oddball and standard stimuli were switched (Figure 3C,D) confirming that the statistics of the stimulus (i.e. rare occurrence of oddball tones versus frequent occurrence of standard tone) drove the larger neural responses to oddball tones rather than tone frequency. The median [IQR] values for phasic and sustained clusters after reversal were 0.32 [0.27, 0.37] and 0.53 [0.38, 0.70] seconds respectively, i.e., the difference in median peak response times (~0.21 seconds) was unchanged following tone reversal.

### Relative timing of phasic and sustained responses to oddball tones

As shown above, phasic cluster responses reliably peaked before sustained cluster responses. To further explore these differences in peak times (Supplementary Figure 3A,B) between the phasic and sustained groups, we used standard cross-correlation analysis to investigate the relative lead or lag between two clusters on a on a trial-by-trial basis. For oddball trials, the phasic cluster responses preceded sustained cluster responses by about 0.14 seconds (Figure 4, red traces). The responses to standard tones did not show any measurable correlations (Figure 4, black traces). We illustrate the relative timing of responses of the two clusters and their anatomical locations by plotting the locations of electrodes in three groups, determined as terciles of the peak time distribution (Figure 5). This reveals a systematic posterior (early; Figure 5A) to anterior (late; Figure 5C) shift that is consistent with early responses in sensorimotor regions and later responses aggregating in salience and frontoparietal regions.

**Figure 4.**
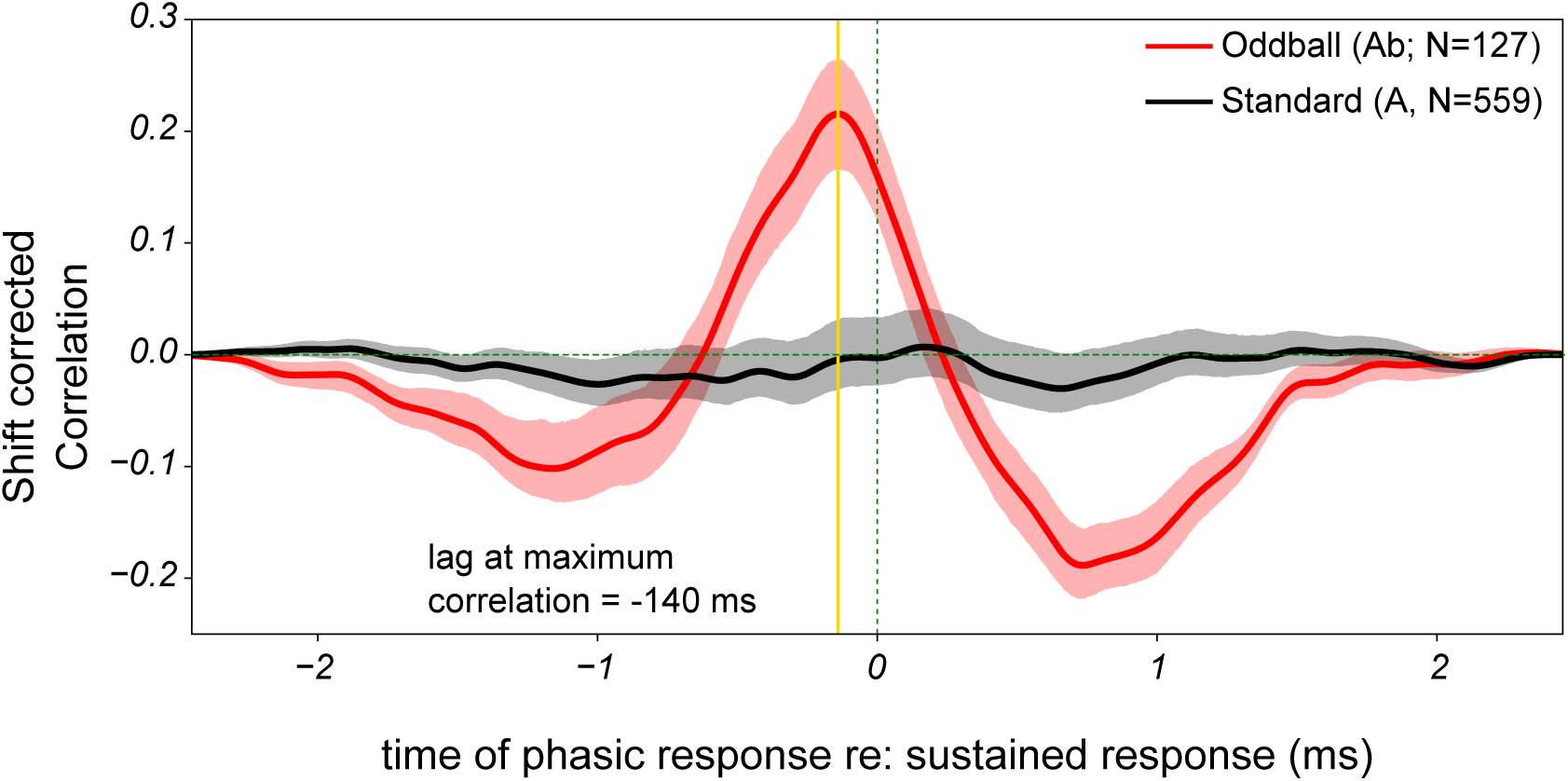
Temporal relationship between phasic and sustained clusters. Cross correlation between phasic and sustained responses for oddball (red) and standard (black) stimuli. The phasic response precedes the sustained by 140 ms. Shaded regions show estimated 95% confidence intervals. The lag at maximum correlation is indicated by the solid vertical line.

**Figure 5.**
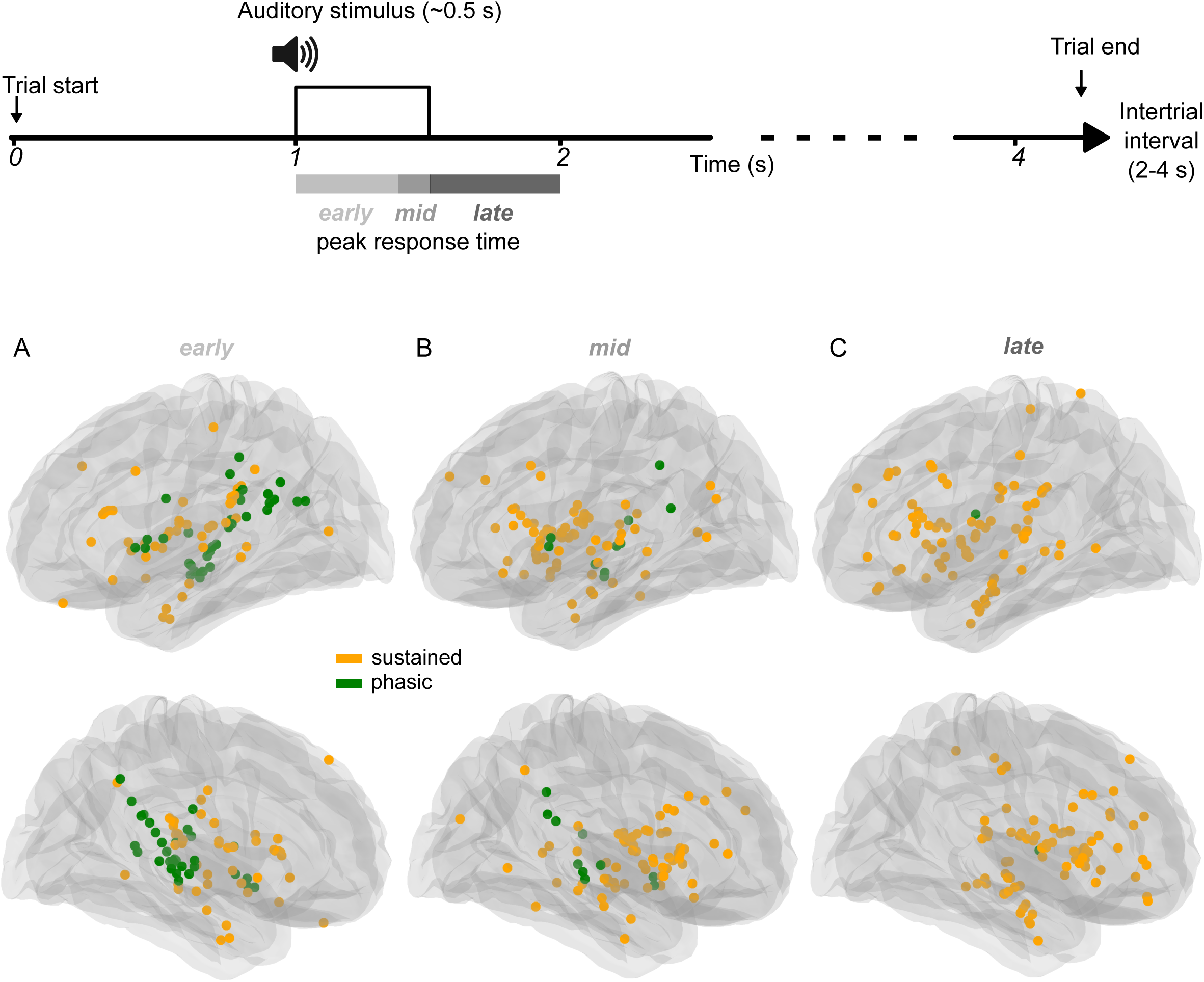
Relative timing of responses in phasic and sustained clusters. Anatomical locations for fast, medium and slow peak response times, calculated as terciles of the peak response time distribution. Task timeline and peak response terciles indicated in top panel. (A) Tercile 1 (0-0.37 s after tone). (B) Tercile 2 (0.37-0.52 s after tone). (C) Tercile 3 (0.52-0.99 s after tone).

## Discussion

We investigated the spatiotemporal dynamics of neural activations evoked by salient auditory stimuli (oddball tones) using high-frequency activity measured using intracranial electrophysiology in neurosurgical patients. We found that: (1) salient auditory stimuli evoked larger neural activations than those evoked by common tones in 22% of the electrodes in two distinct stages: first, a phasic response that peaked relatively early following the oddball tone and returned to baseline rapidly, and second, a sustained response that peaked later and remained elevated for a longer time after tone onset; (2) salience-driven phasic activations, that were distinct from sensory responses, preferentially occurred in sensorimotor regions, whereas sustained activations were largely localized to the salience network; (3) phasic responses reliably preceded sustained responses on a trial-by-trial basis. Taken together, our findings support an updated view of salience processing in the human brain, revealing that salient stimuli evoke two sequential stages of neural activation — phasic sensorimotor responses followed by sustained salience network activity — rather than simultaneous widespread activation.

### Distinct response patterns and timing in different brain networks

Our findings of responses to salient auditory stimuli in the sensorimotor, salience and frontoparietal networks are broadly consistent with single neuron recordings in the macaque (*Camalier et al., 2019*) and EEG or fMRI studies in humans (*Halgren et al., 1998; Yoshiura et al., 1999; Stevens et al., 2000; Liebenthal et al., 2003; Bryant et al., 2005; Sridharan et al., 2007; Watkins et al., 2007*). Similar to our findings, one analysis showed that dorsal attention networks have less of a role in processing rare stimuli (*Kim 2014*). However, the differential distribution of early phasic and later sustained responses among different networks is less well characterized. In one mouse study using visual oddball stimuli, phasic activations in sensory regions evolved over time into sustained activations in integrative regions and circuits (Khilkevich et al., 2024). Event related potential (ERP) studies of responses to salient auditory stimuli have shown early sensory and late cognitive components (*Näätänen et al, 1978; Halgren et al., 1998*) but how these are associated with single unit activity or HFA remains to be determined. Our results might help interpret some of the classic ERP temporal dynamics in terms of the underlying neural activity within functional brain networks.

Our findings are novel because they provide a comprehensive insight into brainwide spatiotemporal dynamics. Previous animal studies of salient auditory stimulus processing used single unit recording to show a range of temporal dynamics in neural responses and have provided some of the motivation for our own efforts. However, they were also limited because they targeted a few brain regions that are thought to be involved in encoding salient events in general or auditory signals in particular (*Paller et al, 1992; Rajkowski et al., 1994; Swick et al., 1994; Ulanovsky et al., 2003; Eliades et al, 2014; Chen et al., 2015; Takaura and Fujii, 2016; Joshi et al, 2016; Camalier et al., 2019; Joshi and Gold, 2022; Gong et al., 2024*). Previous human studies that also used high spatiotemporal resolution intracranial recordings have similar limitations as the animal work. Rather than brainwide recording or analyses, they focused on one or a few brain regions and some only analyzed lower frequency band activity which does not reflect the temporal dynamics of evoked multiunit spiking activity like the HFA that we calculated (*Paller et al, 1992; Zaehle et al, 2013; Citherlet et al, 2019, 2020; Ferraro et al., 2020; Mocchi et al, 2024*). Therefore, while revealing features of neural responses to salient auditory stimuli in specific brain regions, most previous animal and human studies did not describe brainwide networks for salience processing.

### Detection of statistical irregularities – a common cortical and subcortical computation?

The brain’s model of the world, which is an estimate of our environment, is created from statistical regularities of sensory signals experienced by an individual (*Kliuchko et al., 2019; Ho et al., 2022*). In the context of neural computations, prediction errors are the mismatch between the brain’s model of the world and deviations from this model in the brain’s current state (*Rao and Ballard, 1999; Den Ouden et al., 2012*). The detection and encoding of statistical irregularities (like oddball tones) in sensory inputs as prediction errors feeds into key computations in the brain’s circuits. These computations include attentional reorienting and updating the current model of the world and adapting ongoing behaviors that rely on such sensory inputs, resulting in more optimal outcomes (Posner, 1980; Wilson et al., 2010; Nassar et al., 2010; Feldman and Friston, 2010). Such computations are consistent with theories and models of reinforcement learning and adaptive behavior, and with experimental explorations of how and where these signals are encoded (Schultz et al., 1997; Dayan and Daw, 2008; Cohen et al, 2012; Matias et al, 2017).

What mechanisms might generate the selective activations to salient stimuli that we measured? Studies of oddball responses that were influenced by prior scalp EEG measurements (mismatch negativity or MMN; *Näätänen et al, 1978*) showed that the enhanced responses to rare stimuli were due to a cortical, but not thalamic, mechanism called stimulus specific adaptation (*Ulanovsky et al., 2003*). This form of neural adaptation is to the statistical distribution of stimuli: responses to frequent/standard tones adapt while responses to rare tones do not. As a result, rare stimuli evoke relatively large responses compared with frequent ones. Increased tone frequency selective inhibition recruited by frequent stimuli might also contribute to relatively larger responses to oddball stimuli (Valdés-Baizabal et al., 2021). There is evidence that this relative enhancement of the response to rare tones is not purely local but is further amplified by a cortical mechanism involving feedback from higher to lower cortical regions (Fishman and Steinschneider, 2012; Obara et al, 2023). These findings are consistent with theoretical frameworks for optimal neural coding that incorporate the statistical nature of natural sensory environments (*Nelken et al., 1999; Fairhall et al., 2001; Willmore and King, 2022*).

What mechanisms might explain the anatomical separation of phasic and sustained activations to salient stimuli that we observed? One possibility is that salience signals are computed in specific cortical circuits that have descending projections to brainstem nuclei that then broadcast them, via widespread neuromodulator release, to the rest of the brain, perhaps resulting in the delayed, relatively widespread, sustained responses that we measured. This is consistent with the temporal-anatomical pattern of activation that we report here (Figure 5). Phasic salience signals might thus act as prediction error signals, triggering the brainwide neuromodulator release that has been hypothesized to result in a “network reset” (Bouret and Sara, 2005). This process of feedback regulation serves as a reorienting signal by influencing neural processing in downstream circuits, an idea for which evidence has recently started to accumulate (Joshi and Gold, 2022; Grimm et al., 2024). At the circuit level, responses to salient stimuli can influence downstream circuits to modulate ongoing information processing through a range of mechanisms. Bottom-up salient signals gated via neuromodulatory brainstem nuclei provide a pathway for sympathetic regulation of brainwide circuits (Yu and Dayan, 2005; Joshi and Gold, 2020; Cazettes et al, 2021). How these signals are computed remains an open question.

Ours and previous studies have reported the detection and representation of statistical irregularities in sensory stimuli in cortex. How are these findings related with the measurement of enhanced responses to salient auditory stimuli in brainstem structures like the inferior colliculus and locus coeruleus (*Swick et al., 1994; Joshi et al, 2016; Krebs et al, 2018; Chot et al., 2020; Carbajal et al., 2024*)? One explanation is that this is not in fact an exclusively cortical function and that adaptation to repeated stimuli (and consequently, relative enhancement of responses to rare stimuli) is a feature of neural circuits throughout the brain. If the statistical irregularity is complex and multimodal (i.e. a combination of visual, auditory and cognitive factors, among others), then it is less likely to be a brainstem computation as it requires the integration of multiple factors – a computation generally attributed to circuits anchored in cortical hubs (but including thalamic and brainstem regions; *Thiebaut de Schotten and Forkel, 2022*). In this case, brainstem activations are more likely the result of descending projections from integrative cortical regions (*Joshi and Gold, 2020; Salvi et al, 2025*).

### Sparse representation of oddball tones

Many previous studies (see above) using oddball paradigms focused their measurements or analyses on specific brain regions. In our study, we included recordings from all of the locations that had been targeted for monitoring in the epilepsy monitoring unit and found that enhanced responses to oddball tones were relatively sparse (263 of 1194 recording locations). We acknowledge that some brain regions are underrepresented in our data set, particularly motor and occipital cortices, and oddball tones might evoke responses here as well. However, we do not believe that these gaps in coverage could negatively affect our conclusion about sparse representation of auditory salience since a wide range of regions including those in networks associated with salience processing, the focus of our study, were adequately represented in our data.

### Prevalence of phasic and sustained responses in the brain

We showed that responses to oddball tones fell into two broad categories: phasic and sustained. In general, phasic and sustained responses to a range of stimuli and during a variety of behaviors have been measured widely in the brain. Both responses are commonly found in early sensory regions (*Cleland et al., 1971; Williams and Shapley, 2007; Khilkevich et al, 2024*) and are a characteristic feature of salient stimulus evoked neural responses in brainstem, thalamus and cortex (*Swick et al., 1994; Boehnke et al., 2011; Joshi et al., 2016*). Sustained responses in integrative cortical and subcortical regions are often associated with cognitive functions supporting memory processes and evidence accumulation during active behaviors (*Fuster and Alexander, 1971; Fuster et al., 1982; Glimcher and Sparks, 1992; Gold and Shadlen, 2000; Ratcliff et al, 2003*). Responses to salient auditory stimuli, including the types we report here, likely support adaptive, reorienting behaviors by redirecting attention following changes in the environment (Fairhall et al., 2001; Bouret and Sara, 2005; Wilson et al., 2010).

### Salience and brain dysfunction

Understanding salience encoding in the brain also has deep clinical relevance. Individuals suffering from brain injury or neuropsychiatric disorders show deviations in salience processing. For example, in individuals with attention deficit hyperactivity disorder (ADHD), salient stimuli drive activity in brain circuits and sensory modalities that are unrelated to ongoing behavior or task demands and reduced adaptation to familiar stimuli, accompanied by abnormal behavioral responses to irrelevant stimuli (*Stevens et al., 2007; Tegelbeckers et al., 2015; Salmi et al., 2018*). Aberrant neural temporal dynamics and erroneous identification of irrelevant stimuli are signatures of the pathophysiology and cognitive dysfunction in schizophrenia (*Kapur, 2003; Liddle et al., 2006; Wolf et al., 2008; Miyata, 2019; Dewan et al., 2024; Menon et al., 2023*). Similar deficits in salience processing are associated with aging related mild cognitive impairment (MCI; *Staffen et al., 2012; Kurt et al., 2014; Bell et al., 2021*) and traumatic brain injury (TBI; *Boshra et al., 2020; Tapper et al, 2022*). Elucidating the neural dynamics and circuits for salience processing will help us better understand injury and disease driven changes in neural circuits and the accompanying deficits in perception and behavior.

### Limitations

Our study has several limitations. First, it is possible that some of these results are particular to our patient population. In general, epilepsy patients can show additional forms of inter-individual variability in their brain networks related to their pathology (Bettus et al, 2008) that might also manifest in certain categorical behavioral differences compared to healthy controls (Bruhn and Parsons, 1977). However, we sought to mitigate such concerns by focusing on neural signals associated with specific, highly controlled stimuli. Under these kinds of conditions, it has been shown that neural findings from intracranial EEG studies in patients with epilepsy can generalize to healthy controls populations (Long et al., 2014). Nevertheless, more work is needed to fully understand the relationship between behavioral and neural variability across individuals (Genon et al., 2022), which can have broad evolutionary (Bechara et al., 2000), developmental (Tenebaum et al., 2011), and functional (Yang et al., 2020) causes. Second, our clustering analysis was intended to identify distinct functional neural population that were evident across all subjects we studied. A more granular clustering of distinct functional profiles may be possible with a larger dataset.

## Conclusions

Our measurements show that auditory statistical irregularities are reliably encoded in distinct temporal patterns of neural activation within distinct functional brain networks. Specific neural, molecular and circuit mechanisms that generate the response profiles and regional specificity reported here will require further study. Our results lay the groundwork for future studies to determine the functional significance of these clusters for normal brain functions such as learning and attentional reorienting that are central to daily adaptive behaviors. Our work also motivates further explorations of changes in neurocognitive function that follow changes in these networks as a result of traumatic brain injury, neurodegenerative disease, or aging.

## Acknowledgements

We thank Dr. Josef Parvizi for supporting our research at the Stanford EMU, Jiaqi Wu and Creence Lin for their expert help in developing experiment task code, and Zoe Lusk and Ritika Garg for expert help in collecting and curating iEEG data.

---

**Figure 2 Supplement.**
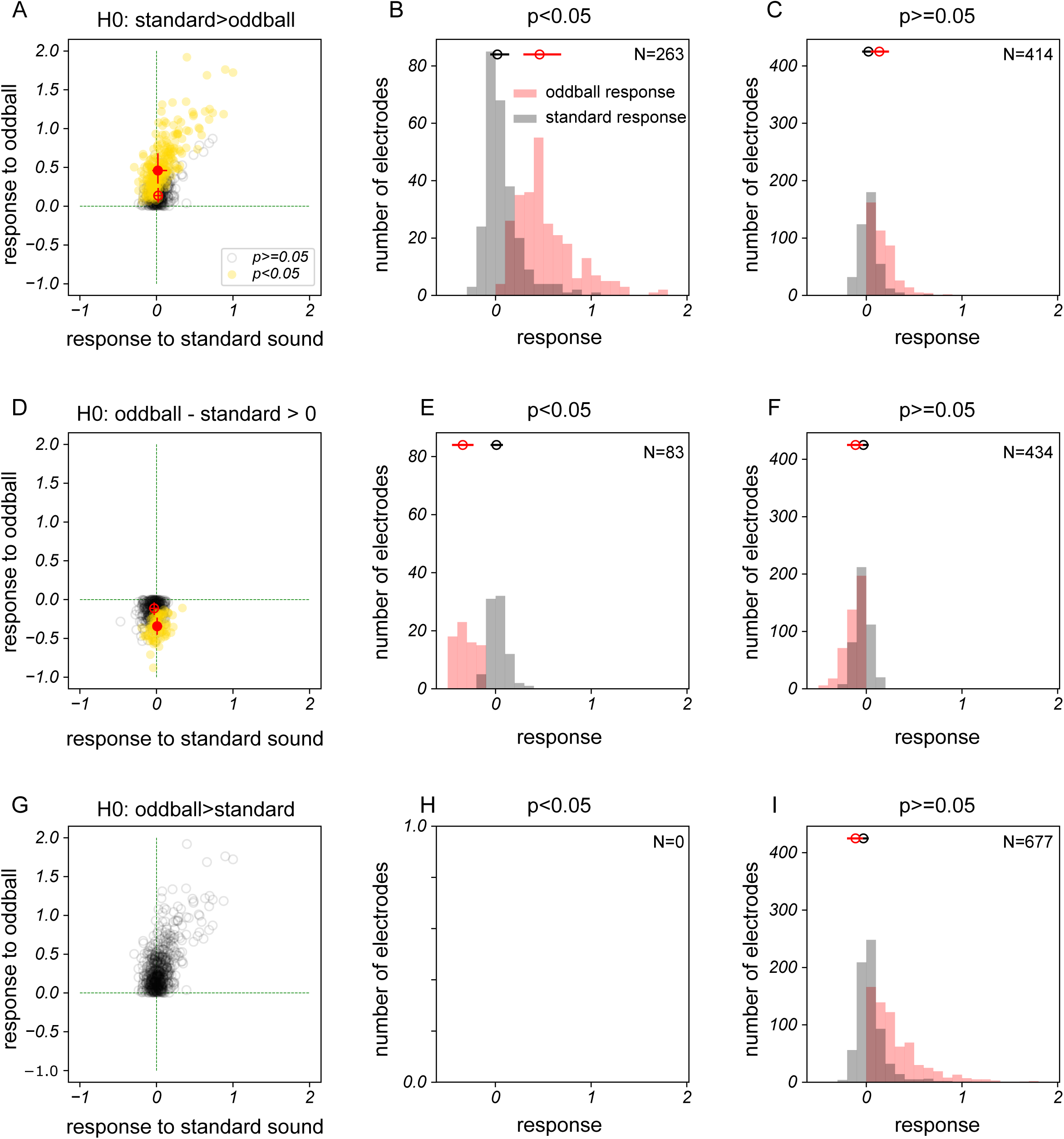
(A) Responses to standard and oddball stimuli. Yellow markers indicate p<0.05, Mann-Whitney U-test for *H0*: response to standard tone > response to oddball tone. Green markers and lines indicate median and interquartile range. (B) and (C) Distributions of the data shown in (A) with markers and lines indicating median and interquartile range. (D-F) Same as (A-C) for cases when the oddball tone suppressed the HFA. (G-I) Same as above for cases where the standard tone evoked a greater response than the oddball tone.

**Figure 3 Supplement.**
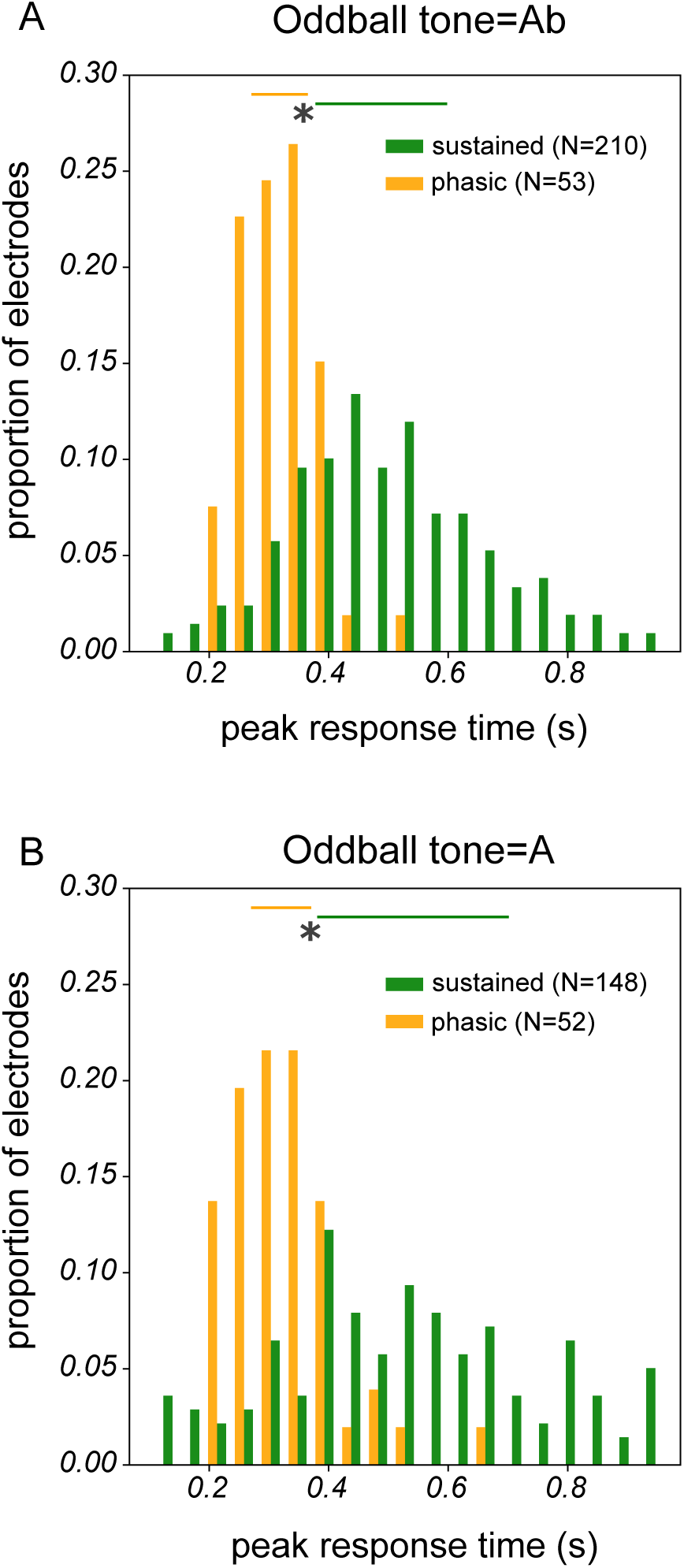
(A) Distribution of peak response time for phasic and sustained clusters. Horizontal bars indicate interquartile range. ‘*’ indicates p<0.005, Mann-Whitney U-test for *H0*: sustained peak time > phasic peak time. (B) Same as (A) but after contingency reversal.

## Notes

### Competing Interest Statement

The authors have declared no competing interest.

### Summary of Updates

We have updated the author list to include the following co-author: Ritika Garg.

### Funding

- National Institutes of Health: R21NS140918-01

## References

1. Bechara A, Damasio H, Damasio AR. (2000). Emotion, decision making and the orbitofrontal cortex. Cereb. Cortex 10: 295–307. doi: 10.1093/cercor/10.3.295. PMID: 10731224.

2. Bettus G, Wendling F, Guye M, Valton L, Régis J, Chauvel P, Bartolomei F. (2008). Enhanced EEG functional connectivity in mesial temporal lobe epilepsy. Epilepsy Res. 81: 58–68. doi: 10.1016/j.eplepsyres.2008.04.020. PMID: 18547787.

3. Boehnke SE, Berg DJ, Marino RA, Baldi PF, Itti L, Munoz DP. (2011). Visual adaptation and novelty responses in the superior colliculus. Eur J Neurosci. 34:766–79. doi: 10.1111/j.1460-9568.2011.07805.x. PMID: 21864319; PMCID: PMC3168683.

4. Boshra R, Ruiter KI, Dhindsa K, Sonnadara R, Reilly JP, Connolly JF. (2020). On the time-course of functional connectivity: theory of a dynamic progression of concussion effects. Brain Commun. 2(2):fcaa063. doi: 10.1093/braincomms/fcaa063. PMID: 32954320; PMCID: PMC7491441.

5. Bouret S, Sara SJ. (2005). Network reset: a simplified overarching theory of locus coeruleus noradrenaline function. Trends Neurosci. 28:574–582. doi: 10.1016/j.tins.2005.09.002. PMID: 16165227.

6. Brázdil M, Dobsík M, Mikl M, Hlustík P, Daniel P, Pazourková M, Krupa P, Rektor I. (2005). Combined event-related fMRI and intracerebral ERP study of an auditory oddball task. Neuroimage. 26:285–293. doi: 10.1016/j.neuroimage.2005.01.051. PMID: 15862229.

7. Bruhn P and Parsons OA. (1977). Reaction time variability in epileptic and brain-damaged patients. Cortex 13: 373–384.

8. Bryant RA, Felmingham KL, Kemp AH, Barton M, Peduto AS, Rennie C, Gordon E, Williams LM. (2005). Neural networks of information processing in posttraumatic stress disorder: a functional magnetic resonance imaging study. Biol Psychiatry. 58:111–118. doi: 10.1016/j.biopsych.2005.03.021. PMID: 16038681.

9. Camalier CR, Scarim K, Mishkin M, Averbeck BB. (2019). A Comparison of Auditory Oddball Responses in Dorsolateral Prefrontal Cortex, Basolateral Amygdala, and Auditory Cortex of Macaque. J Cogn Neurosci. 31:1054–1064. doi: 10.1162/jocn_a_01387. PMID: 30883292; PMCID: PMC6995248.

10. Carbajal GV, Casado-Román L, Malmierca MS. (2024). Two Prediction Error Systems in the Nonlemniscal Inferior Colliculus: “Spectral” and “Nonspectral”. J Neurosci. 44(23):e1420232024. doi: 10.1523/JNEUROSCI.1420-23.2024. PMID: 38627089; PMCID: PMC11154860.

11. Cazettes F, Reato D, Morais JP, Renart A, Mainen ZF. (2021). Phasic Activation of Dorsal Raphe Serotonergic Neurons Increases Pupil Size. Curr Biol. 31:192–197.e4. doi: 10.1016/j.cub.2020.09.090. PMID: 33186549; PMCID: PMC7808753.

12. Chen IW, Helmchen F, Lütcke H. (2015). Specific Early and Late Oddball-Evoked Responses in Excitatory and Inhibitory Neurons of Mouse Auditory Cortex. J Neurosci. 35:12560–12573. doi: 10.1523/JNEUROSCI.2240-15.2015. PMID: 26354921; PMCID: PMC6605391.

13. Chot MG, Tran S, Zhang H. (2020). Spatial Separation between Two Sounds of an Oddball Paradigm Affects Responses of Neurons in the Rat’s Inferior Colliculus to the Sounds. Neuroscience. 444:118–135. doi: 10.1016/j.neuroscience.2020.07.027. PMID: 32712224.

14. Citherlet D, Boucher O, Tremblay J, Robert M, Gallagher A, Bouthillier A, Lepore F, Nguyen DK. (2019). Role of the insula in top-down processing: an intracranial EEG study using a visual oddball detection paradigm. Brain Struct Funct. 224:2045–2059. doi: 10.1007/s00429-019-01892-y. PMID: 31129871.

15. Citherlet D, Boucher O, Tremblay J, Robert M, Gallagher A, Bouthillier A, Lepore F, Nguyen DK. (2020). Spatiotemporal dynamics of auditory information processing in the insular cortex: an intracranial EEG study using an oddball paradigm. Brain Struct Funct. 225:1537–1559. doi: 10.1007/s00429-020-02072-z. PMID: 32347366.

16. Cleland BG, Dubin MW, Levick WR. Sustained and transient neurones in the cat’s retina and lateral geniculate nucleus. J Physiol. 1971 Sep;217:473–496. doi: 10.1113/jphysiol.1971.sp009581. PMID: 5097609; PMCID: PMC1331787.

17. Cohen JD, Aston-Jones G. (2005). Cognitive neuroscience: decision amid uncertainty. Nature. 436(7050):471–472. doi: 10.1038/436471a. PMID: 16049461.

18. Cohen JY, Haesler S, Vong L, Lowell BB, Uchida N. (2012). Neuron-type-specific signals for reward and punishment in the ventral tegmental area. Nature. 482:85–88. doi: 10.1038/nature10754. PMID: 22258508; PMCID: PMC3271183.

19. Critchley HD, Nagai Y, Gray MA, Mathias CJ. (2011). Dissecting axes of autonomic control in humans: Insights from neuroimaging. Auton Neurosci. 161:34–42. doi: 10.1016/j.autneu.2010.09.005. PMID: 20926356.

20. Dayan P, Daw ND. (2008). Decision theory, reinforcement learning, and the brain. Cogn Affect Behav Neurosci. 8:429–453. doi: 10.3758/CABN.8.4.429. PMID: 19033240.

21. Den Ouden HE, Kok P, de Lange FP. (2012). How prediction errors shape perception, attention, and motivation. Front Psychol. 3:548. doi: 10.3389/fpsyg.2012.00548. PMID: 23248610; PMCID: PMC3518876.

22. Devilbiss DM, Waterhouse BD. (2011). Phasic and tonic patterns of locus coeruleus output differentially modulate sensory network function in the awake rat. J Neurophysiol. 105:69–87. doi: 10.1152/jn.00445.2010. PMID: 20980542; PMCID: PMC3023368.

23. Dewan M, Campbell Daniels E, Hunt JE, Bryant EA, Trikeriotis SI, Kelly DL, Adams HA, Hare SM, Waltz JA. (2024). Aberrant salience signaling in auditory processing in schizophrenia: Evidence for abnormalities in both sensory processing and emotional reactivity. Schizophr Res. 274:329–336. doi: 10.1016/j.schres.2024.09.026. PMID: 39454324; PMCID: PMC11620929.

24. Dubey, A., Ray, S. (2019). Cortical Electrocorticogram (ECoG) Is a Local Signal. J. Neurosci. 39: 4299–4311. doi: 10.1523/JNEUROSCI.2917-18.2019. PMID: 30914446; PMCID: PMC6538865.

25. Eliades SJ, Crone NE, Anderson WS, Ramadoss D, Lenz FA, Boatman-Reich D. (2014). Adaptation of high-gamma responses in human auditory association cortex. J Neurophysiol. 112:2147–2163. doi: 10.1152/jn.00207.2014. PMID: 25122702; PMCID: PMC4274926.

26. Fairhall AL, Lewen GD, Bialek W, de Ruyter Van Steveninck RR. (2001). Efficiency and ambiguity in an adaptive neural code. Nature. 412:787–792. doi: 10.1038/35090500. PMID: 11518957.

27. Feldman H, Friston KJ. (2010). Attention, uncertainty, and free-energy. Front Hum Neurosci. 4:215. doi: 10.3389/fnhum.2010.00215. PMID: 21160551; PMCID: PMC3001758.

28. Ferraro S, Van Ackeren MJ, Mai R, Tassi L, Cardinale F, Nigri A, Bruzzone MG, D’Incerti L, Hartmann T, Weisz N, Collignon O. (2020). Stereotactic electroencephalography in humans reveals multisensory signal in early visual and auditory cortices. Cortex. 126:253–264. doi: 10.1016/j.cortex.2019.12.032. PMID: 32092494.

29. Fishman YI, Steinschneider M. (2012). Searching for the mismatch negativity in primary auditory cortex of the awake monkey: deviance detection or stimulus specific adaptation? J Neurosci. 32:15747–15758. doi: 10.1523/JNEUROSCI.2835-12.2012. PMID: 23136414; PMCID: PMC3641775.

30. Fuster JM, Alexander GE. (1971). Neuron activity related to short-term memory. Science. 173:652–4. doi: 10.1126/science.173.3997.652. PMID: 4998337.

31. Fuster JM, Bauer RH, Jervey JP. Cellular discharge in the dorsolateral prefrontal cortex of the monkey in cognitive tasks. Exp Neurol. 1982 Sep;77:679–94. doi: 10.1016/0014-4886(82)90238-2. PMID: 7117470.

32. Genon S, Eickhoff SB, Kharabian S. (2022). Linking interindividual variability in brain structure to behaviour. Nat. Rev. Neurosci. 23: 307–318.

33. Glimcher PW, Sparks DL. (1992). Movement selection in advance of action in the superior colliculus. Nature. 355:542–545. doi: 10.1038/355542a0. PMID: 1741032.

34. Gold JI, Shadlen MN. Representation of a perceptual decision in developing oculomotor commands. Nature. 2000 Mar 23;404:390–394. doi: 10.1038/35006062. PMID: 10746726.

35. Gong Y, Song P, Du X, Zhai Y, Xu H, Ye H, Bao X, Huang Q, Tu Z, Chen P, Zhao X, Pérez-González D, Malmierca MS, Yu X. (2024). Neural correlates of novelty detection in the primary auditory cortex of behaving monkeys. Cell Rep. 43(3):113864. doi: 10.1016/j.celrep.2024.113864. PMID: 38421870.

36. Greicius MD, Krasnow B, Reiss AL, Menon V. (2003). Functional connectivity in the resting brain: a network analysis of the default mode hypothesis. Proc Natl Acad Sci USA. 100:253–258. doi: 10.1073/pnas.0135058100. PMID: 12506194; PMCID: PMC140943.

37. Grimm C, Duss SN, Privitera M, Munn BR, Karalis N, Frässle S, Wilhelm M, Patriarchi T, Razansky D, Wenderoth N, Shine JM, Bohacek J, Zerbi V. (2024). Tonic and burst-like locus coeruleus stimulation distinctly shift network activity across the cortical hierarchy. Nat Neurosci. 27:2167–2177. doi: 10.1038/s41593-024-01755-8. PMID: 39284964; PMCID: PMC11537968.

38. Halgren E, Marinkovic K, Chauvel P. (1998). Generators of the late cognitive potentials in auditory and visual oddball tasks. Electroencephalogr Clin Neurophysiol.106:156–164. doi: 10.1016/s0013-4694(97)00119-3. PMID: 9741777.

39. He H, Hong L, Sajda P. (2023). Pupillary response is associated with the reset and switching of functional brain networks during salience processing. PLoS Comput Biol. 19(5):e1011081. doi: 10.1371/journal.pcbi.1011081. PMID: 37172067; PMCID: PMC10208478.

40. Hillyard SA, Squires KC, Bauer JW, Lindsay PH. (1971). Evoked potential correlates of auditory signal detection. Science. 172:1357–1360. doi: 10.1126/science.172.3990.1357. PMID: 5580218.

41. Ho HT, Burr DC, Alais D, Morrone MC. (2022). Propagation and update of auditory perceptual priors through alpha and theta rhythms. Eur J Neurosci. 55:3083–3099. doi: 10.1111/ejn.15141. PMID: 33559266; PMCID: PMC9543013.

42. Javitt DC, Steinschneider M, Schroeder CE, Vaughan HG Jr, Arezzo JC. (1994). Detection of stimulus deviance within primate primary auditory cortex: intracortical mechanisms of mismatch negativity (MMN) generation. Brain Res. 667:192–200. doi: 10.1016/0006-8993(94)91496-6. PMID: 7697356.

43. Joshi S, Li Y, Kalwani RM, Gold JI. (2016). Relationships between Pupil Diameter and Neuronal Activity in the Locus Coeruleus, Colliculi, and Cingulate Cortex. Neuron. 89:221–234. doi: 10.1016/j.neuron.2015.11.028. PMID: 26711118; PMCID: PMC4707070.

44. Joshi S, Gold JI. (2020). Pupil Size as a Window on Neural Substrates of Cognition. Trends Cogn Sci. Jun;24:466–480. doi: 10.1016/j.tics.2020.03.005. PMID: 32331857; PMCID: PMC7271902.

45. Joshi S, Gold JI. (2022). Context-dependent relationships between locus coeruleus firing patterns and coordinated neural activity in the anterior cingulate cortex. Elife. 11:e63490. doi: 10.7554/eLife.63490. PMID: 34994344; PMCID: PMC8765756.

46. Kapur S. (2003). Psychosis as a state of aberrant salience: a framework linking biology, phenomenology, and pharmacology in schizophrenia. Am J Psychiatry. 160:13–23. doi: 10.1176/appi.ajp.160.1.13. PMID: 12505794.

47. Keil J, Pomper U, Senkowski D. (2016). Distinct patterns of local oscillatory activity and functional connectivity underlie intersensory attention and temporal prediction. Cortex. 74:277–288. doi: 10.1016/j.cortex.2015.10.023. PMID: 26716405.

48. Kerzel D, Schönhammer J. (2013). Salient stimuli capture attention and action. Atten Percept Psychophys. 75(8):1633–1643. doi: 10.3758/s13414-013-0512-3. PMID: 23918550.

49. Khilkevich A, Lohse M, Low R, Orsolic I, Bozic T, Windmill P, Mrsic-Flogel TD. (2024). Brain-wide dynamics linking sensation to action during decision-making. Nature. 634(8035):890–900. doi: 10.1038/s41586-024-07908-w. PMID: 39261727; PMCID: PMC11499283.

50. Kim H. (2014). Involvement of the dorsal and ventral attention networks in oddball stimulus processing: a meta-analysis. Hum Brain Mapp. 35:2265–2284. doi: 10.1002/hbm.22326. PMID: 23900833; PMCID: PMC6868981.

51. Kliuchko M, Brattico E, Gold BP, Tervaniemi M, Bogert B, Toiviainen P, Vuust P. (2019). Fractionating auditory priors: A neural dissociation between active and passive experience of musical sounds. PLoS One. 14(5):e0216499. doi: 10.1371/journal.pone.0216499. PMID: 31051008; PMCID: PMC6499420.

52. Krebs RM, Park HRP, Bombeke K, Boehler CN. (2018). Modulation of locus coeruleus activity by novel oddball stimuli. Brain Imaging Behav. 12(2):577–584. doi: 10.1007/s11682-017-9700-4. PMID: 28271441.

53. Kurt P, Emek-Savaş DD, Batum K, Turp B, Güntekin B, Karşıdağ S, Yener GG. (2014). Patients with mild cognitive impairment display reduced auditory event-related delta oscillatory responses. Behav Neurol. 2014:268967. doi: 10.1155/2014/268967. PMID: 24825953; PMCID: PMC4006610.

54. Lee GR, Gommers R, Wasilewski F, Wohlfahrt K, O’Leary A. (2019). PyWavelets: A Python package for wavelet analysis. Journal of Open Source Software, 4(36), 1237, 10.21105/joss.01237.

55. Liddle PF, Laurens KR, Kiehl KA, Ngan ET. (2006). Abnormal function of the brain system supporting motivated attention in medicated patients with schizophrenia: an fMRI study. Psychol Med. 36:1097–1108. doi: 10.1017/S0033291706007677. PMID: 16650349.

56. Liebenthal E, Ellingson ML, Spanaki MV, Prieto TE, Ropella KM, Binder JR. (2003). Simultaneous ERP and fMRI of the auditory cortex in a passive oddball paradigm. Neuroimage. 19:1395–1404. doi: 10.1016/s1053-8119(03)00228-3. PMID: 12948697.

57. Linden DE, Prvulovic D, Formisano E, Völlinger M, Zanella FE, Goebel R, Dierks T. (1999). The functional neuroanatomy of target detection: an fMRI study of visual and auditory oddball tasks. Cereb Cortex. 9:815–823. doi: 10.1093/cercor/9.8.815. PMID: 10601000.

58. Liu Q, Ulloa A, Horwitz B. (2022). The Spatiotemporal Neural Dynamics of Intersensory Attention Capture of Salient Stimuli: A Large-Scale Auditory-Visual Modeling Study. Front Comput Neurosci. 16:876652. doi: 10.3389/fncom.2022.876652. PMID: 35645750; PMCID: PMC9133449.

59. Long NM, Burke JF and Kahana MJ. (2014). Subsequent memory effect in intracranial and scalp EEG. Neuroimage 84: 488–494.

60. Manning JR, Jacobs J, Fried I, Kahana MJ. (2009). Broadbandshifts in local field potential power spectra are correlated with single-neuron spiking in humans. J. Neurosci. 29, 13613–13620.

61. Matias S, Lottem E, Dugué GP, Mainen ZF. (2017). Activity patterns of serotonin neurons underlying cognitive flexibility. Elife.6:e20552. doi: 10.7554/eLife.20552. PMID: 28322190; PMCID: PMC5360447.

62. Menon V, Palaniyappan L, Supekar K. (2023). Integrative Brain Network and Salience Models of Psychopathology and Cognitive Dysfunction in Schizophrenia. Biol Psychiatry. 94:108–120. doi: 10.1016/j.biopsych.2022.09.029. PMID: 36702660.

63. Miyata J. (2019). Toward integrated understanding of salience in psychosis. Neurobiol Dis. 131:104414. doi: 10.1016/j.nbd.2019.03.002. PMID: 30849509.

64. Mocchi M, Bartoli E, Magnotti J, de Gee JW, Metzger B, Pascuzzi B, Mathura R, Pulapaka S, Goodman W, Sheth S, McGinley MJ, Bijanki K. (2024). Aperiodic spectral slope tracks the effects of brain state on saliency responses in the human auditory cortex. Sci Rep. 14(1):30751. doi: 10.1038/s41598-024-80911-3. PMID: 39730513; PMCID: PMC11681213.

65. Murphy PR, Robertson IH, Balsters JH, O’connell RG. (2011). Pupillometry and P3 index the locus coeruleus-noradrenergic arousal function in humans. Psychophysiology. 48:1532–1543. doi: 10.1111/j.1469-8986.2011.01226.x.. PMID: 21762458.

66. Murphy PR, O’Connell RG, O’Sullivan M, Robertson IH, Balsters JH. Pupil diameter covaries with BOLD activity in human locus coeruleus. Hum Brain Mapp. 2014 Aug;35(8):4140–54. doi: 10.1002/hbm.22466. PMID: 24510607; PMCID: PMC6869043.

67. Näätänen R, Gaillard AW, Mäntysalo S. (1978). Early selective-attention effect on evoked potential reinterpreted. Acta Psychol (Amst). 42(4):313–29. doi: 10.1016/0001-6918(78)90006-9. PMID: 685709.

68. Nassar MR, Wilson RC, Heasly B, Gold JI. (2010). An approximately Bayesian delta-rule model explains the dynamics of belief updating in a changing environment. J Neurosci. 30:12366–12378. doi: 10.1523/JNEUROSCI.0822-10.2010. PMID: 20844132; PMCID: PMC2945906.

69. Nelken I, Rotman Y, Bar Yosef O. (1999). Responses of auditory-cortex neurons to structural features of natural sounds. Nature. 397:154–157. doi: 10.1038/16456. PMID: 9923676.

70. Novembre G, Lacal I, Benusiglio D, Quarta E, Schito A, Grasso S, Caratelli L, Caminiti R, Mayer AB, Iannetti GD. (2024). A Cortical Mechanism Linking Saliency Detection and Motor Reactivity in Rhesus Monkeys. J Neurosci. 44:e0422232023. doi: 10.1523/JNEUROSCI.0422-23.2023. PMID: 37949654; PMCID: PMC10851684.

71. Obara K, Ebina T, Terada SI, Uka T, Komatsu M, Takaji M, Watakabe A, Kobayashi K, Masamizu Y, Mizukami H, Yamamori T, Kasai K, Matsuzaki M. (2023). Change detection in the primate auditory cortex through feedback of prediction error signals. Nat Commun. 14(1):6981. doi: 10.1038/s41467-023-42553-3. PMID: 37957168; PMCID: PMC10643402.

72. Paller KA, McCarthy G, Roessler E, Allison T, Wood CC. (1992). Potentials evoked in human and monkey medial temporal lobe during auditory and visual oddball paradigms. Electroencephalogr Clin Neurophysiol. 84:269–279. doi: 10.1016/0168-5597(92)90008-y. PMID: 1375886.

73. Parasuraman R, Beatty J. (1980). Brain events underlying detection and recognition of weak sensory signals. Science. 210(4465):80–3. doi: 10.1126/science.7414324. PMID: 7414324.

74. Patterson JC 2nd, Ungerleider LG, Bandettini PA. (2002). Task-independent functional brain activity correlation with skin conductance changes: an fMRI study. Neuroimage. 17:1797–1806. doi: 10.1006/nimg.2002.1306. PMID: 12498753.

75. Posner MI. (1980). Orienting of attention. Q J Exp Psychol. Feb;32(1):3–25. doi: 10.1080/00335558008248231. PMID: 7367577.

76. Rajkowski J, Kubiak P, Aston-Jones G. (1994). Locus coeruleus activity in monkey: phasic and tonic changes are associated with altered vigilance. Brain Res Bull. 35:607–616. doi: 10.1016/0361-9230(94)90175-9. PMID: 7859118.

77. Rao RP, Ballard DH. (1999). Predictive coding in the visual cortex: a functional interpretation of some extra-classical receptive-field effects. Nat Neurosci. 2(1):79–87. doi: 10.1038/4580. PMID: 10195184.

78. Ratcliff R, Cherian A, Segraves M. (2003). A comparison of macaque behavior and superior colliculus neuronal activity to predictions from models of two-choice decisions. J Neurophysiol. 90:1392–407. doi: 10.1152/jn.01049.2002. PMID: 12761282.

79. Ray S, Maunsell JH. (2011). Different origins of gamma rhythm and high-gamma activity in macaque visual cortex. PLoS Biol. 9, e1000610.

80. Salmi J, Salmela V, Salo E, Mikkola K, Leppämäki S, Tani P, Hokkanen L, Laasonen M, Numminen J, Alho K. (2018). Out of focus - Brain attention control deficits in adult ADHD. Brain Res. 1692:12–22. doi: 10.1016/j.brainres.2018.04.019. PMID: 29702087.

81. Salvi V, Courtand G, de Deurwaerdère P, Cardoit L, Valerio S, Delcasso S, Georges F, Michelet T. (2025). Cingulate cortex stimulation drives distinct pupillary responses in rat via recruitment of noradrenergic neurons in the locus coeruleus. Cereb Cortex. 35(5):bhaf085. doi: 10.1093/cercor/bhaf085. PMID: 40315430.

82. Schultz W, Dayan P, Montague PR. (1997). A neural substrate of prediction and reward. Science. 275(5306):1593–9. doi: 10.1126/science.275.5306.1593. PMID: 9054347.

83. Seeley WW, Menon V, Schatzberg AF, Keller J, Glover GH, Kenna H, Reiss AL, Greicius MD. (2007). Dissociable intrinsic connectivity networks for salience processing and executive control. J Neurosci. 27:2349–2356. doi: 10.1523/JNEUROSCI.5587-06.2007. PMID: 17329432; PMCID: PMC2680293.

84. Seeley, W. W. (2019). The salience network: a neural system for perceiving and responding to homeostatic demands. Journal of Neuroscience, 39: 9878–9882. doi: 10.1523/JNEUROSCI.1138-17.2019. PMID: 31676604; PMCID: PMC6978945.

85. Sridharan D, Levitin DJ, Chafe CH, Berger J, Menon V. (2007). Neural dynamics of event segmentation in music: converging evidence for dissociable ventral and dorsal networks. Neuron. 55:521–532. doi: 10.1016/j.neuron.2007.07.003. PMID: 17678862.

86. Staffen W, Ladurner G, Höller Y, Bergmann J, Aichhorn M, Golaszewski S, Kronbichler M. (2012). Brain activation disturbance for target detection in patients with mild cognitive impairment: an fMRI study. Neurobiol Aging. 33(5):1002.e1–16. doi: 10.1016/j.neurobiolaging.2011.09.002. PMID: 21993055.

87. Stevens AA, Skudlarski P, Gatenby JC, Gore JC. (2000). Event-related fMRI of auditory and visual oddball tasks. Magn Reson Imaging. 18(5):495–502. doi: 10.1016/s0730-725x(00)00128-4. PMID: 10913710.

88. Stevens MC, Pearlson GD, Kiehl KA. (2007). An FMRI auditory oddball study of combined-subtype attention deficit hyperactivity disorder. Am J Psychiatry. 164(11):1737–1749. doi: 10.1176/appi.ajp.2007.06050876. PMID: 17974940.

89. Sutton RS, Barto AG. (1998). Reinforcement Learning: An Introduction Cambridge, MA: MIT Press.

90. Swick D, Pineda JA, Schacher S, Foote SL. (1994). Locus coeruleus neuronal activity in awake monkeys: relationship to auditory P300-like potentials and spontaneous EEG. Exp Brain Res. 101:86–92. doi: 10.1007/BF00243219. PMID: 7843306.

91. Takaura K, Fujii N. (2016). Facilitative effect of repetitive presentation of one stimulus on cortical responses to other stimuli in macaque monkeys--a possible neural mechanism for mismatch negativity. Eur J Neurosci. 43:516–528. doi: 10.1111/ejn.13136. PMID: 26613160; PMCID: PMC5064748.

92. Tapper A, Staines WR, Niechwiej-Szwedo E. (2022). EEG reveals deficits in sensory gating and cognitive processing in asymptomatic adults with a history of concussion. Brain Inj. 36:1266–1279. doi: 10.1080/02699052.2022.2120210. PMID: 36071612.

93. Tegelbeckers J, Bunzeck N, Duzel E, Bonath B, Flechtner HH, Krauel K. (2015). Altered salience processing in attention deficit hyperactivity disorder. Hum Brain Mapp. 36:2049–2060. doi: 10.1002/hbm.22755. PMID: 25648705; PMCID: PMC4670482.

94. Tenenbaum JB, Kemp C, Griffiths TL, Goodman ND. (2011). How to grow a mind: statistics, structure, and abstraction. Science 331: 1279–1285. doi: 10.1126/science.1192788. PMID: 21393536.

95. Thiebaut de Schotten M, Forkel SJ. (2022). The emergent properties of the connected brain. Science. 378(6619):505–510. doi: 10.1126/science.abq2591. PMID: 36378968.

96. Uddin LQ, Yeo BTT, Spreng RN. (2019). Towards a Universal Taxonomy of Macro-scale Functional Human Brain Networks. Brain Topogr. 32(6):926–942. doi: 10.1007/s10548-019-00744-6. PMID: 31707621; PMCID: PMC7325607.

97. Ulanovsky N, Las L, Nelken I. (2003). Processing of low-probability sounds by cortical neurons. Nat Neurosci. 2003 6:391–398. doi: 10.1038/nn1032. PMID: 12652303.

98. Usher M, Cohen JD, Servan-Schreiber D, Rajkowski J, Aston-Jones G. (1999). The role of locus coeruleus in the regulation of cognitive performance. Science. 283:549–54. doi: 10.1126/science.283.5401.549. PMID: 9915705.

99. Valdés-Baizabal C, Casado-Román L, Bartlett EL, Malmierca MS. (2021). In vivo whole-cell recordings of stimulus-specific adaptation in the inferior colliculus. Hear Res. 399:107978. doi: 10.1016/j.heares.2020.107978. PMID: 32402412.

100. Wang CA, Boehnke SE, Itti L, Munoz DP. (2014) Transient pupil response is modulated by contrast-based saliency. J Neurosci. 34:408–417. doi: 10.1523/JNEUROSCI.3550-13.2014. PMID: 24403141; PMCID: PMC6608151.

101. Watkins S, Dalton P, Lavie N, Rees G. (2007). Brain mechanisms mediating auditory attentional capture in humans. Cereb Cortex. 17:1694–1700. doi: 10.1093/cercor/bhl080. PMID: 16990437.

102. Williams PE, Shapley RM. (2007). A dynamic nonlinearity and spatial phase specificity in macaque V1 neurons. J Neurosci. 27:5706–5718. doi: 10.1523/JNEUROSCI.4743-06.2007. PMID: 17522315; PMCID: PMC6672757.

103. Willmore BDB, King AJ. (2023). Adaptation in auditory processing. Physiol Rev. 103:1025–1058. doi: 10.1152/physrev.00011. PMID: 36049112; PMCID: PMC9829473.

104. Wilson MJ, Harkrider AW, King KA. (2014). The effects of visual distracter complexity on auditory evoked p3b in contact sports athletes. Dev Neuropsychol. 39:113–130. doi: 10.1080/87565641.2013.870177. PMID: 24571930.

105. Wilson RC, Nassar MR, Gold JI. (2010). Bayesian online learning of the hazard rate in change-point problems. Neural Comput. 22:2452–2476. doi: 10.1162/NECO_a_00007. PMID: 20569174; PMCID: PMC2966286.

106. Wolf DH, Turetsky BI, Loughead J, Elliott MA, Pratiwadi R, Gur RE, Gur RC. (2008). Auditory Oddball fMRI in Schizophrenia: Association of Negative Symptoms with Regional Hypoactivation to Novel Distractors. Brain Imaging Behav. 2:132–145. doi: 10.1007/s11682-008-9022-7. PMID: 19756228; PMCID: PMC2743436.

107. Yang, G. R. & Wang, X. J. Artificial neural networks for neuroscientists: a primer. Neuron 107, 1048–1070 (2020). doi: 10.1016/j.neuron.2021.01.022. PMID: 32970997; PMCID: PMC11576090.

108. Yeo BT, Krienen FM, Sepulcre J, Sabuncu MR, Lashkari D, Hollinshead M, Roffman JL, Smoller JW, Zöllei L, Polimeni JR, Fischl B, Liu H, Buckner RL. (2011). The organization of the human cerebral cortex estimated by intrinsic functional connectivity. J Neurophysiol. 106:1125–65. doi: 10.1152/jn.00338.a2011. PMID: 21653723; PMCID: PMC3174820.

109. Yoshiura T, Zhong J, Shibata DK, Kwok WE, Shrier DA, Numaguchi Y. (1999). Functional MRI study of auditory and visual oddball tasks. Neuroreport. 10:1683–8. doi: 10.1097/00001756-199906030-00011. PMID: 10501557.

110. Yu AJ, Dayan P. (2005). Uncertainty, neuromodulation, and attention. Neuron. 46:681–692. doi: 10.1016/j.neuron.2005.04.026. PMID: 15944135.

111. Zaehle T, Bauch EM, Hinrichs H, Schmitt FC, Voges J, Heinze HJ, Bunzeck N. (2013). Nucleus accumbens activity dissociates different forms of salience: evidence from human intracranial recordings. J Neurosci. 33:8764–8771. doi: 10.1523/JNEUROSCI.5276-12.2013. PMID: 23678119; PMCID: PMC6618843.

